# Dual proteomics of *Drosophila melanogaster* hemolymph infected with the heritable endosymbiont *Spiroplasma poulsonii*

**DOI:** 10.1101/2021.02.22.432267

**Authors:** Florent Masson, Samuel Rommelaere, Alice Marra, Fanny Schüpfer, Bruno Lemaitre

## Abstract

Insects are frequently infected with heritable bacterial endosymbionts. Endosymbionts have a dramatic impact on their host physiology and evolution. Their tissue distribution is variable with some species being housed intracellularly, some extracellularly and some having a mixed lifestyle. The impact of extracellular endosymbionts on the biofluids they colonize (e.g. insect hemolymph) is however difficult to appreciate because biofluids composition can depend on the contribution of numerous tissues.

Here we address the question of *Drosophila* hemolymph proteome response to the infection with the endosymbiont *Spiroplasma poulsonii. S. poulsonii* inhabits the fly hemolymph and gets vertically transmitted over generations by hijacking the oogenesis in females. Using dual proteomics on infected hemolymph, we uncovered a low level and chronic activation of the Toll immune pathway by *S. poulsonii* that was previously undetected by transcriptomics-based approaches. Using Drosophila genetics, we also identified candidate proteins putatively involved in controlling *S. poulsonii* growth. Last, we also provide a deep proteome of *S. poulsonii*, which, in combination with previously published transcriptomics data, improves our understanding of the post-transcriptional regulations operating in this bacterium.

**Summary statement:** We report the changes in *Drosophila melanogaster* hemolymph proteome upon infection with the heritable bacterial endosymbiont *Spiroplasma poulsonii*.

## Introduction

Insects frequently harbor vertically transmitted bacterial symbionts living within their tissues, called endosymbionts (Douglas, 2015). Endosymbionts have a major impact on the host physiology and evolution as they provide ecological benefits such as the ability for the host to thrive on unbalanced diets (Douglas, 2016), protection against viruses or parasites (Hedges et al., 2008; Oliver et al., 2003; Scarborough et al., 2005; Teixeira et al., 2008) or tolerance to heat (Montllor et al., 2002). A peculiar group of insect endosymbionts also directly affects their host reproduction. This group is taxonomically diverse and includes bacteria from the *Wolbachia, Spiroplasma, Arsenophonus, Cardinium* and *Rickettsia* genera (Duron et al., 2008; Medina et al., 2019). Four reproduction-manipulative mechanisms have been unraveled so far, namely cytoplasmic incompatibility, male-killing, parthenogenesis and male feminization (Werren et al., 2008), all of them leading to an evolutionary drive that favors infected individuals over non-infected ones and promotes the endosymbiont spread into populations. Their ease of spread coupled with the virus-protecting ability of some species (Hedges et al., 2008; Teixeira et al., 2008) make them promising tools to control insect pest populations or to render them refractory to human arboviruses (Hoffmann et al., 2011). Reproductive manipulators are however fastidious bacteria that are difficult to manipulate, hence slowing down research on the functional aspects of their interaction with their hosts (Masson and Lemaitre, 2020). Their real impact on host physiology thus remains largely elusive.

*Spiroplasma* are long helical bacteria belonging to the Mollicutes class, which are devoid of cell wall (Gasparich, 2002). They infect a wide range of arthropods and plants where they act as pathogens, commensals or vertically transmitted endosymbionts depending on the species (Gasparich, 2002). *S. poulsonii* (hereafter *Spiroplasma*) is a vertically transmitted endosymbiont infecting the fruit fly *Drosophila* (Haselkorn, 2010; Mateos et al., 2006). It lives free in the host hemolymph where it feeds on circulating lipids (Herren et al., 2014) and achieves vertical transmission by infecting oocytes (Herren et al., 2013). Most strains cause male-killing, that is the death of the male progeny during early embryogenesis through the action of a toxin named Spaid (Harumoto and Lemaitre, 2018). *Spiroplasma* infection also confers protection to the fly or its larvae against major natural enemies such as parasitoid wasps (Ballinger and Perlman, 2017; Xie et al., 2014) and nematodes (Hamilton et al., 2016; Jaenike et al., 2010).

*Spiroplasma* has long been considered as untractable, but recent technical advances such as the development of *in vitro* culture (Masson et al., 2018) and transformation (Masson et al., 2020b) make it an emergent model for studying the functional aspects of insect-endosymbiont interactions. Some major steps have been made in the understanding of the male killing (Harumoto et al., 2014; Harumoto et al., 2016; Harumoto et al., 2018; Veneti et al., 2005) and protection phenotypes (Ballinger and Perlman, 2017; Hamilton et al., 2016; Paredes et al., 2016) or on the way the bacterial titer was kept under control in the adult hemolymph (Herren et al., 2014; Paredes et al., 2015). Such studies, however, relied on single-gene studies and did not capture the full picture of the impact of *Spiroplasma* on its host physiology. We tackled this question using dual-proteomics on fly hemolymph infected by *Spiroplasma*. This non-targeted approach allowed us to get an extensive overview of the end-effect of *Spiroplasma* infection on the fly hemolymph protein composition and to identify previously unsuspected groups of proteins that where over- or under-represented in infected hemolymph. Using *Drosophila* genetics to knock-down the corresponding genes, we identified new putative regulators of the bacterial titer. This work also provides the most comprehensive *Spiroplasma* proteome to date. By comparing this proteome to the existing transcriptomics data, we draw conclusions about *Spiroplasma* post-transcriptional regulations.

## Results

### 1. *Drosophila* hemolymph protein profiling

We investigated the effect of *Spiroplasma* infection on *Drosophila* hemolymph protein content using Liquid Chromatography-tandem Mass Spectrometry (LC-MS/MS). To this end, we extracted total hemolymph from uninfected and infected 10 days old females. At this age, *Spiroplasma* is already present at high titers in the hemolymph but does not cause major deleterious phenotypes to the fly (Herren and Lemaitre, 2011; Masson et al., 2020a). Extraction was achieved by puncturing the thorax and drawing out with a microinjector. This procedure induces some tissue damage but recovers very few hemocytes (circulating immunes cells). Peptides were mapped to both the *Drosophila* and *Spiroplasma* predicted proteomes, hence allowing having an overview of i) differentially represented *Drosophila* proteins in infected versus uninfected hemolymph and ii) an in-depth *Spiroplasma* proteome in the infected hemolymph samples. These two datasets will be analyzed in separate sections.

The mapping on *Drosophila* proteome allowed the identification of 909 quantified protein groups (a protein group contains proteins that cannot be distinguished based on the identified peptides). The complete list of quantified *Drosophila* proteins and their fold-change upon *Spiroplasma* infection is available in Supplementary Table S1. The hemolymph extraction process involves tissue damage (e.g. the subcuticular epithelium, muscles and fat body) that leads to the release of intracellular proteins in the sample. Hence we first sorted protein groups depending on whether the main protein is predicted to be extracellular or not. Proteins were defined as extracellular if they bore a predicted signal peptide or were annotated with a Gene Ontology (GO) term “extracellular”, or intracellular if no signal peptide nor any “extracellular” GO term was predicted. Based on this localization criteria the *Drosophila* dataset could be split in 555 intracellular protein groups representing 61% of the total proteome and 354 extracellular proteins representing the remaining 39% of the total proteome (Figure 1A).

**Figure 1.**
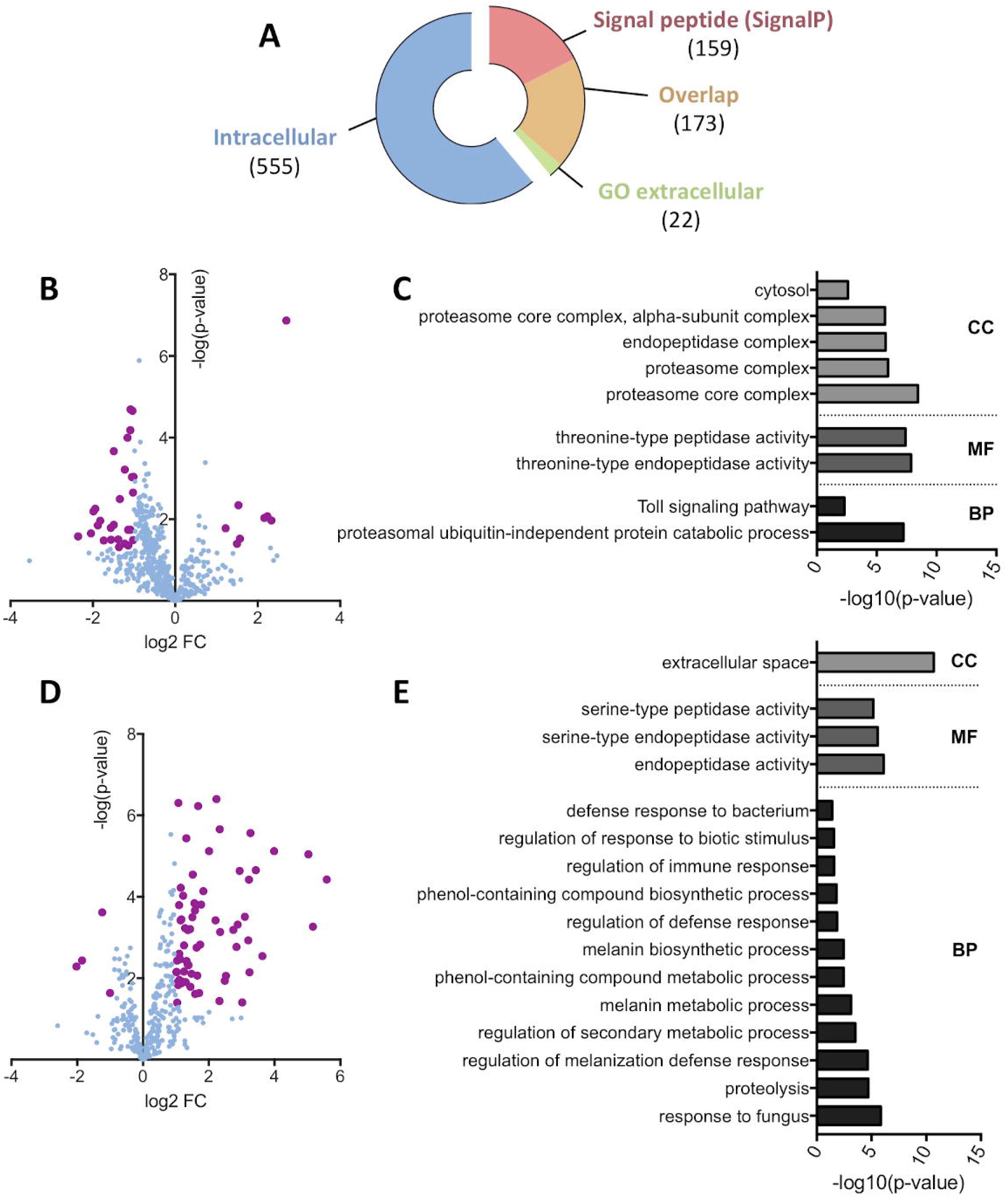
*Drosophila* hemolymph protein profiling upon *Spiroplasma* infection. (A) Representation of the dataset repartition between intracellular and extracellular protein groups. Extracellular groups are detected either by the presence of a predicted signal peptide using the SignalP algorithm or the annotation with a GO term referring to the extracellular space. The overlap category corresponds to protein groups with both a predicted signal peptide and a GO term “extracellular”. Numbers in brackets indicate the number of protein groups in each category. (B) Volcano plot of the intracellular protein group subset. The log fold-change indicates the differential representation of the protein group in the infected samples versus the uninfected samples. Bold purple dots indicate significance as defined by an absolute fold change value over 2 or under -2 and a p-value below 0.05. (C) Functional GO annotation of the intracellular protein groups. Only significant (p-value < 0.05) GO terms of hierarchy 4 or less are displayed. CC: Cell Component; MF: Molecular Function; BP: Biological Process. (D) Volcano plot of the extracellular protein group subset. The log fold-change indicates the differential representation of the protein group in the infected samples versus the uninfected samples. Bold purple dots indicate significance as defined by an absolute fold change value over 2 or under -2 and a p-value below 0.05. (E) Functional GO annotation of the extracellular protein groups. Only significant (p-value < 0.05) GO terms of hierarchy 4 or less are displayed. CC: Cell Component; MF: Molecular Function; BP: Biological Process.

We then compared the relative amounts of protein groups between infected and uninfected hemolymph. Of the intracellular protein groups, 35 were differentially represented in the *Spiroplasma* infected samples, including 8 overrepresented and 27 depleted groups compared to the uninfected control (Figure 1B). A functional GO analysis on the differentially represented protein groups revealed that a significant part of these protein groups were related to the proteasome (subunits α2, 3, 4 and 7 and subunits β3 and 6) and the Toll immune pathway (Figure 1C). The Toll pathway GO enrichment comprises only proteasome subunits, probably because of their involvement in the degradation of Toll pathway intermediates (e.g. Cactus (Palombella et al., 1994)). While differentially abundant intracellular protein groups were mostly underrepresented in *Spiroplasma* infected hemolymph, analysis of the extracellular protein subset revealed an opposite trend. Among the 71 extracellular protein groups having a significantly different abundance, 4 were depleted and 67 were overrepresented in the infected samples compared to the uninfected ones (Figure 1D). Surprisingly, the functional GO annotation highlighted an overrepresentation of serine-endopeptidase molecular function. Serine proteases are involved in the regulation of the melanization and the Toll pathways and indeed the GO terms are enriched for these biological processes as well (Figure 1E).

### 2. Immune-related protein enrichment is a consequence of a mild transcriptional activation

Since *Spiroplasma* is devoid of cell wall, it lacks microbe-associated molecular patterns such as peptidoglycan and was not expected to interact with pattern-recognition receptors regulating the fly immune system. This idea was supported by previous work showing that *Spiroplasma* do not trigger a strong level of Toll and Imd pathway, the two main immune pathways in *Drosophila* (Herren and Lemaitre, 2011; Hurst et al., 2003). Previous work also showed that flies defective for the inducible humoral immune response have normal *Spiroplasma* titer, suggesting that immune pathways are not required to control *Spiroplasma* growth (Herren and Lemaitre, 2011). Our observation that several immune-related proteins are more abundant in the hemolymph of infected flies (Figure 2A and Supplementary Table S1) led us to investigate further on this point. We first tested if the changes in immune-related protein abundance were a consequence of altered gene expression. We measured the corresponding mRNA levels of several differentially represented proteins (Figure 2B). Immune-related genes were slightly upregulated in infected flies, although their induction levels did not compare with those observed after systemic infection by a pathogenic bacteria (De Gregorio et al., 2001; Troha et al., 2018). These results confirm that *Spiroplasma* does not trigger a strong immune response (Herren and Lemaitre, 2011; Hurst et al., 2003). But our data reveals that the presence of *Spiroplama* still provokes a mild and chronic production of immunity proteins.

**Figure 2.**
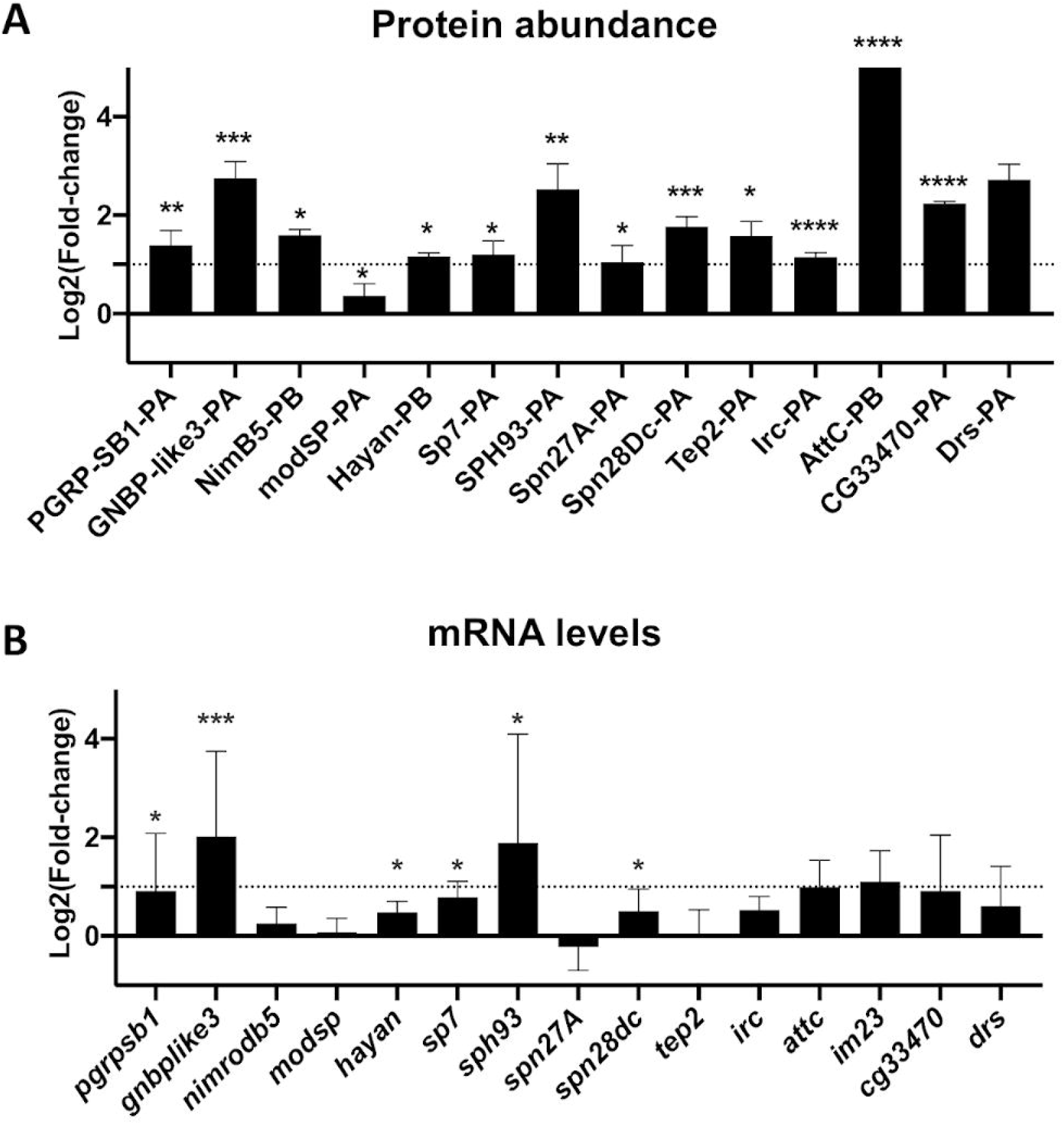
Spiroplasma infection induces a mild transcriptional immune response. (A) Changes in a selection of immune-related protein group abundances quantified by LC-MS/MS. Data are expressed as fold change of the Label-Free Quantification (LFQ) values of *Spiroplasma* infected hemolymph over the uninfected control (log2 scale). *, p<0.05; ** p<0.005, ***; p<0.005, ****, p<0.0005 upon Student t-test. (B) mRNA quantification of a selection of candidate genes in uninfected and *Spiroplasma*-infected flies. Results are expressed as the fold change of target mRNA normalized by *rpL32* mRNA in *Spiroplasma*-infected flies as compared to uninfected flies (log2 scale). *, p<0.05; ** p<0.005, ***; p<0.0005 upon Mann-Whitney test on ΔCt values.

### 3. A majority of enriched hemolymphatic proteins are not involved in *Spiroplasma* growth control

A systemic infection or a genetic induction of the humoral immune response promotes *Spiroplasma* growth in flies (Herren and Lemaitre, 2011). We therefore wondered if the presence of small amounts of antimicrobial peptides (AMPs) could be beneficial for *Spiroplasma* growth. To test this hypothesis, we measured *Spiroplasma* titer in flies over-expressing different AMP genes (Tzou et al., 2002). We found that AMP gene overexpression did not increase *Spiroplasma* titer (Figure 3A). Similarly, flies lacking ten AMP coding genes (Hanson et al., 2019) had a *Spiroplasma* titer comparable to that of control flies (Figure 3B). These results suggest that AMPs released during the humoral immune response do not promote *Spiroplasma* growth.

**Figure 3.**
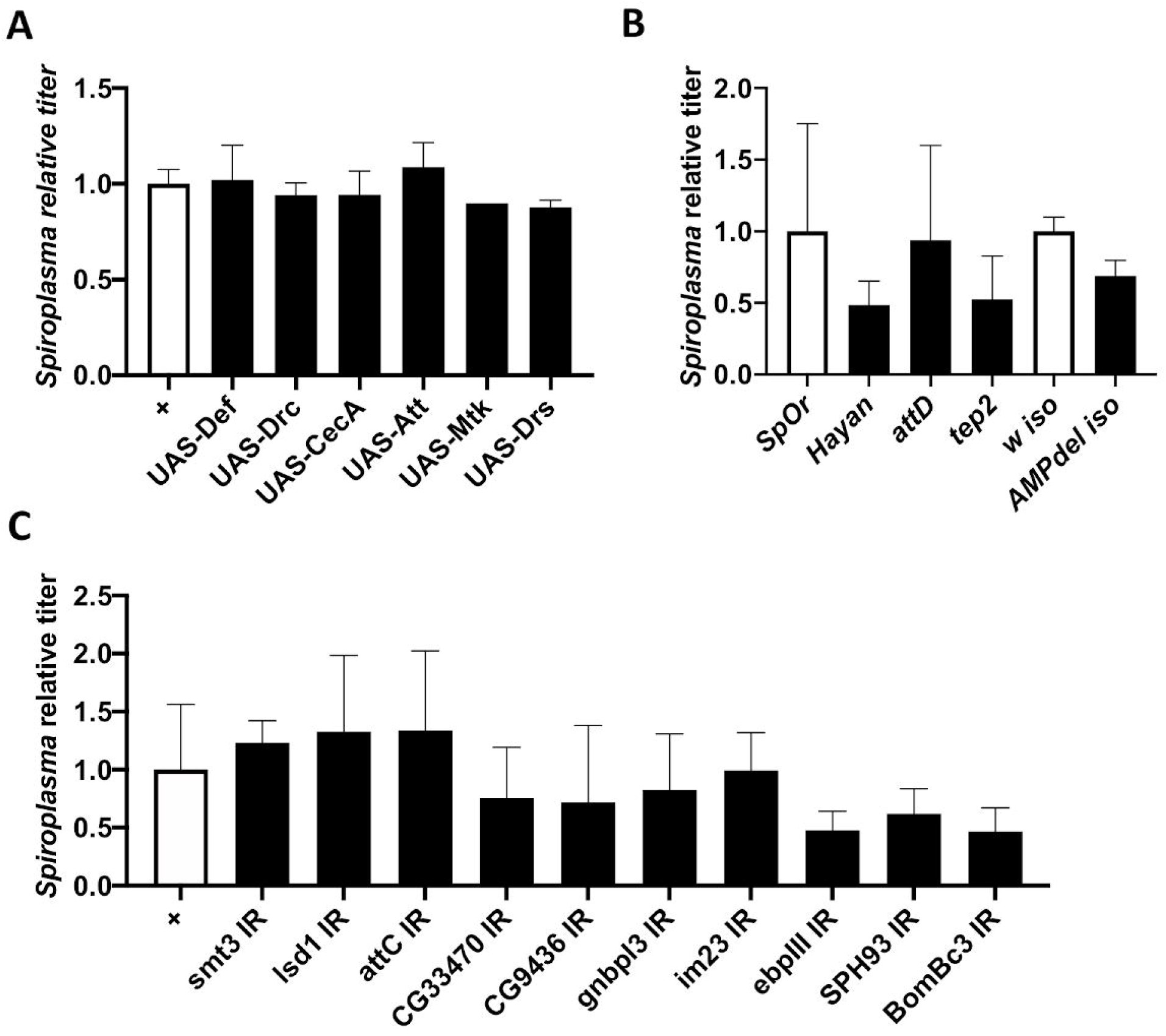
Targeted genetic screening of candidate proteins. (A-C) Quantification of *Spiroplasma* titer in two weeks old flies in various genetic backgounds. Titer is expressed as the fold-change over the appropriate control line. All graphs represent the mean +/-standard deviation of a pool of at least 2 independent experiments. (A) Control flies (Act5C-GAL4 driver crossed w^1118^, white bars) are compared to flies overexpressing a single antimicrobial peptide gene driven by the ubiquitous *Act5C-GAL4* driver (black bars). Titer is expressed as the fold-change over control. (B) Quantification of *Spiroplasma* titer in wild-type flies (*Oregon*^*R*^, white bar) and *hayan, attD* and *tep2* null mutants (black). Spiroplasma titer in AMP10 iso flies were compared to that of their isogenic wild-type counterpart (*w*^*1118*^ iso, white bar). (C) Quantification of the impact of RNAi-mediated knockdown on *Spiroplasma* titer with the Act5C-GAL4 driver. Titer is expressed as the fold-change over the control (Act5C-GAL4 driver crossed to *w*^*1118*^).

We then tested if other immune-related proteins found more abundant in infected hemolymph could alter *Spiroplasma* growth. We first compared the *Spiroplasma* titers of control *Oregon*^*R*^ flies with that of *hayan, attD* and *tep2* mutants. We observed no difference in bacterial titer, indicating that these genes are not required neither detrimental to *Spiroplasma* growth (Figure 3B). We then used an *in vivo* RNAi approach to silence immune genes coding for the most enriched proteins in *Spiroplasma*-infected flies using the ubiquitously expressed *Act5C-GAL4* driver. Silencing these genes did not provoke an increase in *Spiroplasma* titer, further reinforcing the idea that immune proteins do not prevent *Spiroplasma* growth (Figure 3C).

Among the most differentially represented proteins in *Spiroplasma* infected hemolymph, we also detected several proteins of unknown function. To test their role in *Spiroplasma* growth control, we used the same RNAi-mediated knockdown approach and quantified *Spiroplasma* titer in these flies. Most of the RNAi lines tested showed no change in *Spiroplasma* titer as compared to control lines (Supplementary Figure S1). Two RNAi lines targeting *CG18067* and *CG14762*, respectively, showed a slight yet not significant reduction in *Spiroplasma* titer. These results suggest that all the tested genes do not facilitate *Spiroplasma* growth nor control its titer. Instead, these proteins are likely to be induced as a consequence of *Spiroplasma* infection and may participate in the host adaptation to the bacteria. A mutant line was available for only one of the most enriched uncharacterized proteins (*CG15293*). We found that this mutation leads to a significant increase in bacteria titer (Supplementary Figure S1). This gene codes for a 37 kDa secreted protein with no known function. Our results raised the hypothesis that *CG15293* participates in the control of *Spiroplasma* titer *in vivo*.

### 4. Transcriptome-proteome correlation in *Spiroplasma poulsonii*

The proteomics analysis of infected hemolymph samples also allowed the detection and quantification of *Spiroplasma* proteins. With a total of 503 proteins, this is the most comprehensive *Spiroplasma* proteome to date. The complete list of *Spiroplasma* quantified proteins is available in Supplementary Table S2.

We took advantage of a previously published transcriptomics dataset of *Spiroplasma* (Masson et al., 2018) to compare the expression level of *Spiroplasma* genes to their corresponding protein abundance by building a linear model of the proteomics signal as a function of the transcript level (Figure 4). The two measures are poorly correlated (Kendall’s τ = 0.40). Analyzing the normalized residuals of the model also allowed us to identify proteins that are particularly discrepant with the model, that is with absolute standardized residuals greater that 2. This approach uncovered 27 proteins, of which 11 have a significantly higher abundance and 16 a lower abundance in the proteome that what was predicted from the transcriptomics data (Supplementary Table S3). Over-represented proteins with a reliable annotation include the membrane lectin Spiralin B, the cytoskeletal protein Fibrilin, the glucose transporter Crr, the potassium channel KtrA, a ferritin-like protein Ftn, the chromosome partitioning protein ParA and the DNA polymerase subunit DnaN. Under-represented proteins include the transporters SteT and YdcV, the serine/threonine phosphatase Stp, the helicase PcrA, the ATP synthase subunit AtpH, the GMP reductase GuaC and the tRNA-(guanine-N(7)-)-methyltransferase TrmB. Other proteins have no predicted function.

**Figure 4.**
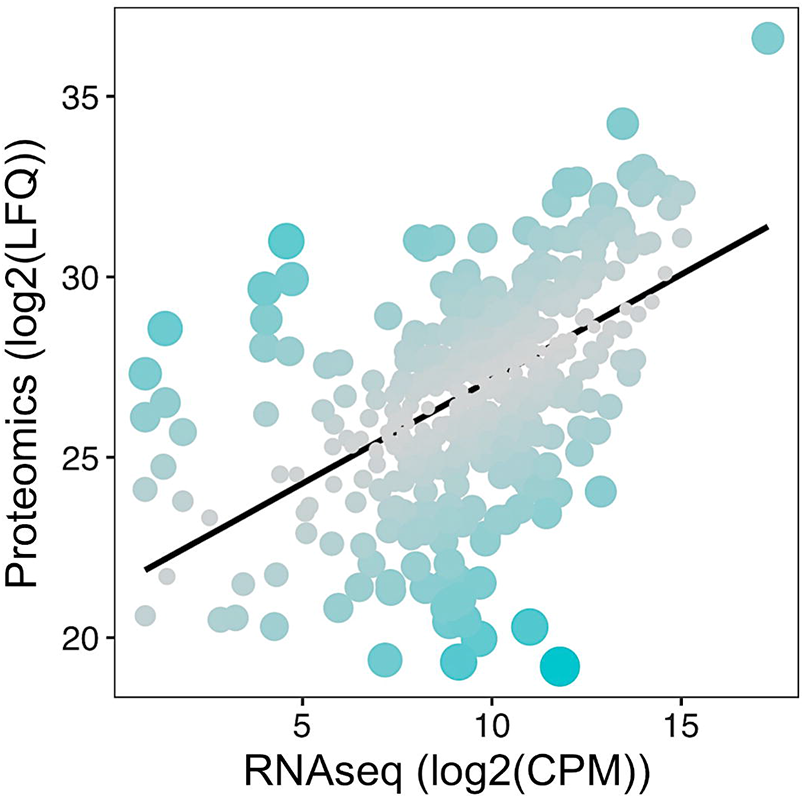
*Spiroplasma* transcription-translation correlation. Each dot represents a protein positioned according to its log2(LFQ; Label-free Quantification) value in the proteome versus its log2(CPM; Count Per Million) in the transcriptomics dataset from (Masson et al., 2018). The black line represents the linear model log2(LFQ) = 0.57892 x log2(CPM) + 21.38325, with an adjusted R^2^ = 0.2909. Dot size and color are adjusted to the residuals of the model, with bigger bluer dots indicating a higher residual hence a stronger deviation from the linear model for the considered protein.

## Discussion

This study provides an extensive characterization of the proteome of *Spiroplasma*-infected *Drosophila* hemolymph. This dual-proteomics approach provides a deep proteome of *Spiroplasma* in its natural environment and pinpoints host proteins which abundance is altered by the presence of the bacteria. The power of *Drosophila* genetics allowed us to test the involvement of these candidate genes in regulating endosymbiosis. A targeted genetic screen revealed that most of the differentially abundant proteins upon *Spiroplasma* infection do not participate in the control of symbiont growth but may rather be involved in host adaptation to *Spiroplasma*.

*Spiroplasma* is devoid of cell wall, hence devoid of peptidoglycan, which is the main immune elicitor for the insect immune system (Lemaitre and Hoffmann, 2007). This led to the assumption that *Spiroplasma* was undetectable by the canonical innate immune pathways. Furthermore, the elicitation of the fly immune system (by an infection or using genetics) has no impact on *Spiroplasma* titer, suggesting that the bacteria are not only invisible by also resistant to the fly immune effectors (Herren and Lemaitre, 2011). Flies that over-express AMPs and the *ΔAMP10* mutant line have normal bacterial titer that confirms the host immune system does not affect *Spiroplasma* growth. However, numerous immune-related proteins (mostly associated to the Toll pathway) were enriched in *Spiroplasma* infected hemolymph, including soluble receptors (GNBP1, GNBP-like3), signal transduction intermediates (Spätzle-Processing Enzyme and Hayan) and effectors or putative effectors (Attacin, Bomanin Bc3 and CG33470). The *Spiroplasma* titer was not altered in several flies carrying loss-of function alleles of these genes, indicating that the immune elicitation is a consequence of the infection but not a control mechanism. It is worth noting that the expression levels of the genes were extremely low as compared to a fully-fledged immune response to pathogenic bacteria (De Gregorio et al., 2001). Such low induction of the immune response has been reported in flies experiencing stress, such as cold or heat stress (MacMillan et al., 2016; Telonis-Scott et al., 2013). Therefore, the mild induction of immune proteins in infected hemolymph may be an unspecific stress response associated to the presence of bacteria. Another attractive hypothesis would be that proteases released by *Spiroplasma* could trigger the soluble sensor Persephone and activate the Toll pathway in a peptidoglycan-independent fashion (Gottar et al., 2006; Issa et al., 2018).

Another uncovering of this study is the depletion of proteasome components in the hemolymph upon *Spiroplasma* infection. As the ubiquitin-proteasome is a major degradation pathway for intracellular proteins (Kleiger and Mayor, 2014), the components that we detected in the hemolymph are presumably released from cells that broke upon hemolymph collection (epithelial, fat or muscular cells, but also possibly hemocytes). However, functionally active 20S proteasome units have been discovered circulating in extracellular fluids in mammals, including in serum (Sixt and Dahlmann, 2008) hence we cannot exclude the existence of circulating proteasome units in *Drosophila* hemolymph. Host ubiquitin-proteasome systems have long been suspected to be a key element in symbiotic homeostasis. It is upregulated in cells harboring intracellular symbionts in weevils, presumably to increase protein turnover and provide free amino-acids to the bacteria (Heddi et al., 2005). *In vitro* work on cell cultures infected by the facultative endosymbiont *Wolbachia* also revealed that it induces the host proteasome presumably also to support its own growth (Fallon and Witthuhn, 2009; He et al., 2019; White et al., 2017). Remarkably, one proteasome subunit was detected as enriched upon *Spiroplasma* infection in the aphid *Aphis citricidus* when the insect was feeding on a suboptimum but not on an optimum host plant, suggesting an interaction between symbiosis, nutrition and the proteasome-ubiquitin degradation system (Guidolin et al., 2018). In the case of *Drosophila-Spiroplasma* interaction, the depletion of proteasome subunits upon infection could thus be a titer regulation mechanism: by decreasing its proteolysis, the host would limit the release of amino acids in the extracellular space that would be usable by *Spiroplasma*.

Our approach also produced an in-depth *Spiroplasma* proteome on total proteins regardless of their cell localization. This gives a quantitatively unbiased overview of each protein abundance which was not achieved by previous data based on detergent extractions (Paredes et al., 2015). Such quantitative approach allowed us to make a comparison between the transcript and protein levels on about one fourth of the total number of predicted coding sequences (Masson et al., 2018). Although the correlation between transcriptomics and proteomics data greatly varies among models, tissues and experimental set-ups (Haider and Pal, 2013), our data indicate a rather low correlation in the case of *Spiroplasma*. The correlations between transcript and protein levels depends on the interaction between transcript stability and protein stability (Schwanhäusser et al., 2011). It also depends on the intrinsic chemical properties of each transcript that affect the ribosome binding or the translation speed, for example the Shine-Dalgarno sequence in prokaryotes (Bingham et al., 1986) or more importantly the codon adaptation index of the coding sequence (Lithwick and Margalit, 2003). An explanation for the overall lack of correlation between transcripts and proteins regardless of the model could be that selection operates at the protein level, hence changes in mRNA levels would be offset by post-transcriptional mechanisms to maintain stable protein levels over evolutionary times (Schrimpf et al., 2009). This hypothesis entails that genes having a low mRNA level compared to their protein levels are likely to be under strong selective pressure to maintain high protein abundancy through post-transcriptional regulations. Intriguingly, this is the case for *S. poulsonii* Spiralin B, a protein homologous to the *S. citri* Spiralin A, a membrane lectin suspected to be essential for insect to plant transmission (Duret et al., 2003; Killiny et al., 2005). Spiralin B has been identified as a putative virulence factor in *S. poulsonii* as it is upregulated when the bacteria are in the fly hemolymph compared to *in vitro* culture (Masson et al., 2018) and could be involved in oocyte infection for vertical transmission. Similarly, the Crr glucose transporter has an unexpectedly high protein/mRNA ratio, possibly in connection with its role in bacteria survival in the hemolymph. *Spiroplasma* has a pseudogenized transporter for trehalose, the most abundant sugar in the hemolymph (Paredes et al., 2015). This could have been selected over host-symbiont coevolution to prevent *Spiroplasma* overgrowth, hence increasing the stability of the interaction by limiting the metabolic cost of harboring the bacteria. Maintaining high Crr levels could thus be an offset response to maintain bacteria proliferation despite trehalose inaccessibility. Last, the ferritin-like protein Ftn has also a bias towards high protein abundancy, suggesting that iron could be a key metabolite in *Spiroplasma*-*Drosophila* symbiotic homeostasis.

All tissues bathed by hemolymph contribute to its composition. As a consequence, studying the impact of symbionts on this biofluid is unapproachable by transcriptomic methods only. Dual proteomics proved to be an efficient approach to overcome this hurdle and gain novel insights into the *Drosophila*-*Spiroplasma* symbiosis. This method is also promising for the study of other symbioses, particularly those where symbionts inhabit biofluids rather than cells.

## Material and Methods

### *Spiroplasma* and *Drosophila* stocks

Flies were kept at 25°C on cornmeal medium (35.28 g of cornmeal, 35.28 g of inactivated yeast, 3.72 g of agar, 36 ml of fruits juice, 2.9 ml of propionic acid and 15.9 ml of Moldex for 600 ml of medium). The complete list of fly stocks is available in Supplementary Table S4.

*Spiroplasma poulsonii* strain Uganda-1 (Pool et al., 2006) was used for all infections. Fly stocks were infected as previously described (Herren and Lemaitre, 2011). Briefly, 9 nL of *Spiroplasma*-infected hemolymph was injected in their thorax. The progeny of these flies was collected after 5-7 days using male killing as a proxy to assess the infection (100% female progeny). Flies from at least the 3^rd^ generation post-injection were used experimentally.

### Hemolymph extraction procedure

Embryos were collected from conventionally reared flies and dechorionated using a bleach-based previously published method (Broderick et al., 2014) to ensure that they were devoid of horizontally transmitted pathogens. One generation was then left to develop in conventional rearing conditions to allow for gut microbiota recontamination. Hemolymph was extracted from 10 days-old flies from the second generation after bleach treatment using a Nanoject II (Drummond) as previously described (Herren et al., 2014). 1 µL of hemolymph was collected for each sample and frozen at -80°C until further use. Hemolymph was then diluted 10 times with PBS containing Protease Inhibitor Cocktail 1X (Roche). 1 µl was used for protein quantification with the Pierce BCA Protein Assay Kit (Thermofisher). The remaining 9 µl were mixed with SDS (0.2% final), DTT (2.5mM final) and PMSF (10µM final). Aliquots of 15µg were used for proteomics analysis.

### Proteomics sample preparation and LC-MS/MS

Each sample was digested by Filter Aided Sample Preparation (FASP) (Wiśniewski et al., 2009) with minor modifications. Dithiothreitol (DTT) was replaced by Tris (2-carboxyethyl)phosphine (TCEP) as reducing agent and iodoacetamide by chloracetamide as alkylating agent. A proteolytic digestion was performed using Endoproteinase Lys-C and Trypsin. Peptides were desalted on C18 StageTips (Rappsilber et al., 2007) and dried down by vacuum centrifugation. For LC-MS/MS analysis, peptides were resuspended and separated by reversed-phase chromatography on a Dionex Ultimate 3000 RSLC nanoUPLC system in-line connected with an Orbitrap Q Exactive HF Mass Spectrometer (Thermo Fischer Scientific). Database search was performed using MaxQuant 1.5.1.2 (Cox and Mann, 2008) against a concatenated database consisting of the Ensemble *Drosophila melanogaster* protein database (BDGP5.25) and a custom *Spiroplama poulsonii* proteome predicted from the reference genome (Genbank accession number JTLV00000000.2). Carbamidomethylation was set as fixed modification, whereas oxidation (M) and acetylation (Protein N-term) were considered as variable modifications. Label-free quantification was performed by MaxQuant using the standard settings. Hits were considered significant when the fold-change between infected and uninfected hemolymph was >2 or <-2 and the FDR<0.05. Complete proteomics data are available on the ProteomExchange repository with the accession number #[data under submission].

### *Spiroplasma* quantification

*Spiroplasma* quantification was performed by qPCR as previously described (Masson et al., 2020a). Briefly, the DNA of pools of 5 whole flies was extracted and the copy number of a single-copy bacterial gene (*dnaA* or *dnaK*) was quantified and normalized by that of the host gene *rsp17*. Primers sequences are available in Supplementary Table S4. Each experiment has been repeated 2 or 3 independent times with at least 3 technical replicates each. Data were analyzed by one-way ANOVA.

### RT-qPCR

Gene expression measurement were performed by RT-qPCR as previously described (Iatsenko et al., 2020; Romeo and Lemaitre, 2008). Briefly, 10 whole flies were crushed and their RNA extracted with the Trizol method. Reverse transcription was carried out using a PrimeScript RT kit (Takara) and a mix of random hexamers and oligo-dTs. qPCR was performed on a QuantStudio 3 (Applied Biosystems) with PowerUp SYBR Green Master Mix using primers sequences available in Supplementary Table S4. The expression of the target gene was normalized by that of the housekeeping gene *rp49* (*rpL32*) using the 2-ΔΔCT method (Pfaffl, 2001). Each experiment has been repeated 2 or 3 independent times with at least 3 technical replicates each. Data were analyzed by Mann-Whitney tests.

## Acknowledgments

We are grateful to Florence Armand, Romain Hamelin and the EPFL Proteomics Core Facility for sharing their expertise on proteomics.

---

**Supplementary Figure 1**.

(A) Quantification of *Spiroplasma* titer in two weeks old flies. Titer is expressed as the fold-change of *Act5C-GAL4>UAS-RNAi* over *Act5C-GAL4> w*^*1118*^. Bars represent the mean +/-standard deviation of a pool of at least 2 independent experiments. (B) Quantification of *Spiroplasma* titer in *CG15293* mutant flies compared to control *yw* flies. Bars represent the mean +/-standard deviation of a pool of at least 3 independent experiments. ***; p<0.0005 upon Mann-Whitney test on ΔΔCt values.

**Supplementary Table 1**.

*Drosophila* genes quantified by LC-MS/MS.

**Supplementary Table 2**.

*Spiroplasma* genes quantified by LC-MS/MS.

**Supplementary Table 3**.

*Spiroplasma* genes outlying the linear model between mRNA and protein levels.

