## Supplementary figures and images for "Dual proteomics of *Drosophila melanogaster* hemolymph infected with the heritable endosymbiont *Spiroplasma poulsonii*"

### Supplementary Figure S1

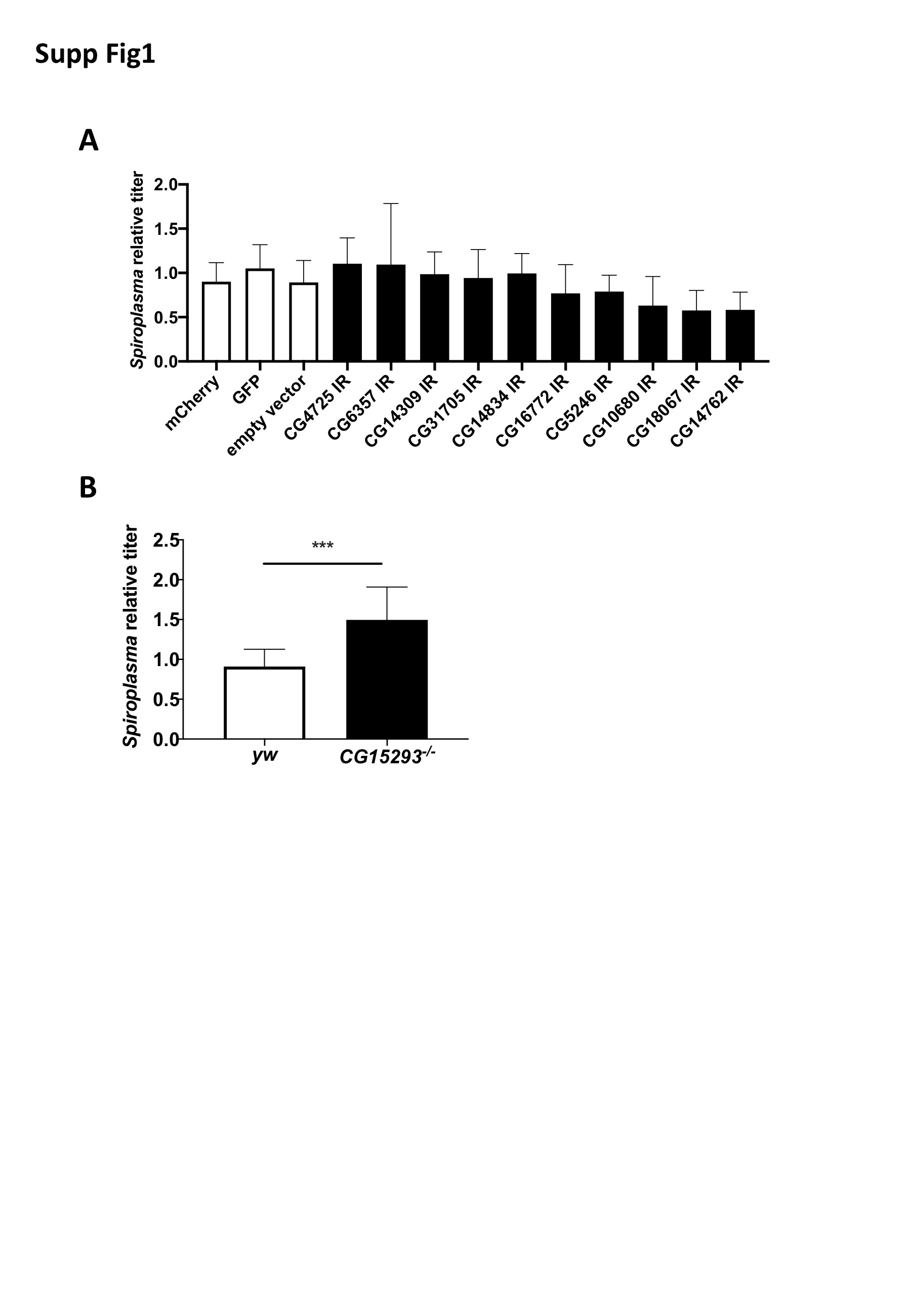
